# A novel method for characterizing cell-cell interactions at single-cell resolution reveals unique signatures in blood T cell-monocyte complexes during infection

**DOI:** 10.1101/2024.09.20.612103

**Authors:** Ningxin Kang, Ashu Chawla, Hannah Hillman, Rashmi Tippalagama, Cheryl Kim, Zbigniew Mikulski, Grégory Seumois, Pandurangan Vijayanand, Thomas J Scriba, Aruna D De Silva, Angel Balmaseda, Eva Harris, Daniela Weiskopf, Alessandro Sette, Cecilia Lindestam Arlehamn, Bjoern Peters, Julie G Burel

**Affiliations:** Center for Vaccine Innovation, La Jolla Institute for Immunology, CA 92037, United States; Bioinformatics Core, La Jolla Institute for Immunology, CA 92037, United States; Flow Cytometry Core, La Jolla Institute for Immunology, CA 92037, United States; Microscopy Core, La Jolla Institute for Immunology, CA 92037, United States; Center for Autoimmunity and Inflammation, La Jolla Institute for Immunology, CA, United States; Department of Medicine, Division of Infectious Diseases and Global Public Health, University of California San Diego (UCSD), La Jolla, CA 92037, USA; South African Tuberculosis Vaccine Initiative (SATVI), Institute of Infectious Disease and Molecular Medicine, Division of Immunology, Department of Pathology, University of Cape Town, South Africa; Faculty of Medicine, General Sir John Kotelawala Defence University, Sri Lanka; Sustainable Sciences Institute, Managua, Nicaragua; Division of Infectious Diseases and Vaccinology, School of Public Health, University of California Berkeley, Berkeley, CA 94720-3370, USA

## Abstract

Communication between immune cells through direct contact is a critical feature of immune responses. Here, we developed a novel high-throughput method to study the transcriptome and adaptive immune receptor repertoire of single cells forming complexes without needing bioinformatic deconvolution. We found that T cells and monocytes forming complexes in blood during active tuberculosis (TB) and dengue hold unique transcriptomic signatures indicative of TCR/MCH-II immune synapses. Additionally, T cells in complexes showed enrichment for effector phenotypes, imaging and transcriptomic features of active TCR signaling, and increased immune activity at diagnosis compared to after anti-TB therapy. We also found evidence for bidirectional RNA exchange between T cells and monocytes, since complexes were markedly enriched for “dual-expressing” cells (i.e., co-expressing T cell and monocyte genes). Thus, studying immune cell complexes at a single-cell resolution offers novel perspectives on immune synaptic interactions occurring in blood during infection.

**Graphical Abstract:** 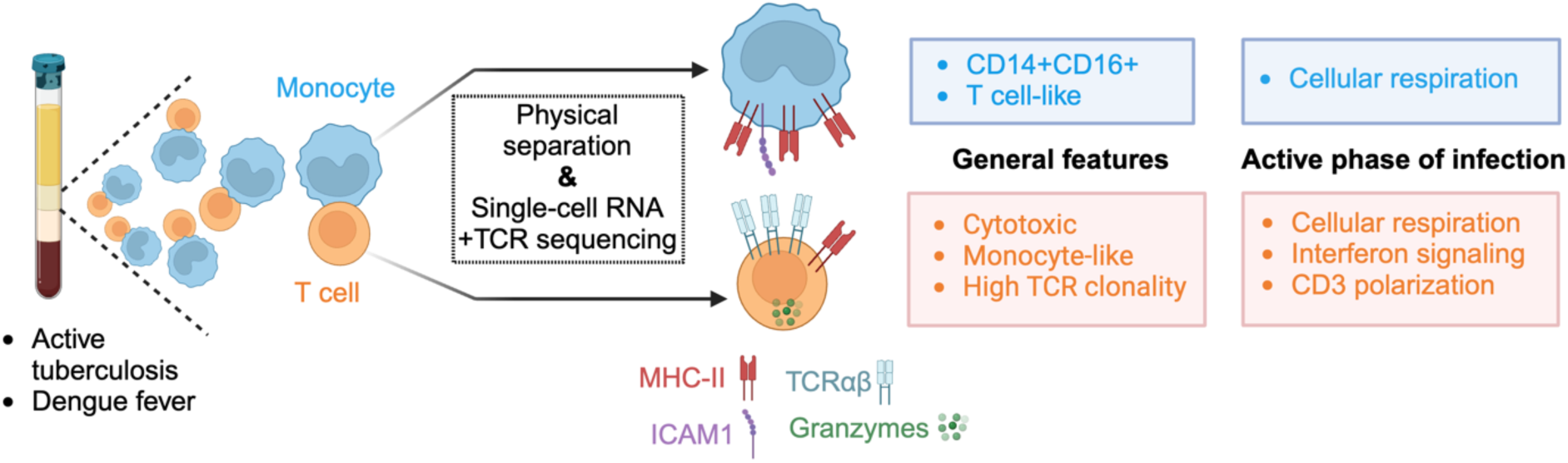

## Introduction

Direct contact between immune cells is a key signaling modality during immune responses. A prototypical example is the immune synapse formation between a T cell and an antigen-presenting cell (APC) through direct interaction between T cell receptor (TCR) and major histocompatibility complex (MHC) molecules (1, 2). During infection, pathogen-derived peptides are processed by APCs and loaded on their MHC at the cell surface. Peptide-MHC complexes are then recognized by T cells expressing a matching TCR on their surface. Upon TCR/peptide-MHC interaction, polarization of CD3, as well as adhesion molecules LFA1 and ICAM1 occurs at the point of contact between the two cells (1, 2). In addition, there is recruitment of other co-stimulatory molecules such as TNFR family members 4-1BB/4-1BBL and OX40/OX40L and rearrangement of cytoskeleton to stabilize the interaction (2, 3). At the transcriptional level, some of the earliest events in T cells following TCR engagement are the activation of Ca2+–calcineurin, mitogen-activated protein kinase (MAPK), and nuclear factor-kB (NFkB) signaling pathways (4). By far the most studied immune synapses are between T cell and B cell engineered cell lines *in vitro*, but there is an increasing understanding that their features likely vary for primary cells *in vivo* and for other APC types such as dendritic cells and monocytes (5).

Our previous work discovered the presence of T cell-monocyte complexes in human blood analyzed by flow cytometry which resemble bona fide biological interactions (6). T cell-monocyte complexes were detected directly from whole blood with minimal sample manipulation, showed LFA1/ICAM1 polarization at their point of contact, and their frequency fluctuated over time following immune perturbations, such as active tuberculosis (ATB), dengue, or Tdap boost vaccination (6). Since then, multiple other groups have described the presence of T cell-monocyte complexes in human blood using either flow cytometry, or single-cell transcriptomics, with increased prevalence in SARS-CoV2 infection (7, 8), cancer (9–11), and chronic inflammation (12–15). For instance, in a high-dimensional flow cytometry analysis of blood from COVID-19 convalescent individuals, two of the three myeloid subsets with increased prevalence compared to healthy controls co-expressed CD3 and CD14 (7). More recently, Carniti et al. elegantly demonstrated that the presence of T cell-monocyte complexes in blood samples of a subset of patients with lymphoma negatively affects the outcome of CAR-T cell therapy (11). So far, all studies have solely focused on associating the frequency of T cell-monocyte complexes with clinical features, and the immune information contained within circulating T cell-monocyte complexes remains uncharacterized.

The study of cell-cell complexes is challenging. We have demonstrated that flow cytometry-derived parameters often fail to identify doublets, resulting in a “contamination” in the singlet cell gate that complicates data interpretation (16). Another major hurdle in the molecular study of cell-cell complexes is that they are detected as one single event by flow cytometers and thus analyzed as a whole, representing a mixture signal from its two cellular components (16). Deconvolution of the signal into each cell component has been elegantly demonstrated as possible at the transcriptomic level, for instance, with the PIC-Seq (17) or ProximID (18) assays. However, this approach is limited to genes that are only expressed by one cell type of the complex (i.e., lineage-specific genes) and precludes the analysis of gene programs that are shared by both cell types, which is the case for the majority of cellular and biological processes.

For this study, we set out to understand the biology of T cell-monocyte complexes in blood, in particular by defining their transcriptomic signatures during infection. We elected ATB as the primary model of study, and dengue as a validation model. Both T cells and monocytes are known to play a role in the immune response to ATB (19–22) and dengue (23, 24), and we have previously identified a higher likelihood of forming complexes at the acute time of infection in both disease models (6). We designed a novel high-throughput method to directly measure the RNA content of individual cells forming complexes, bypassing the need for bioinformatic deconvolution. Together, we identified cellular, protein, gene, and TCR signatures specific to T cells and monocytes forming complexes, furthering our understanding of their mechanisms of adhesion and immune function during infection.

## Results

### Determination of the single-cell transcriptome of cells forming complexes

Thus far, methods for studying cell-cell complexes on a large scale rely on sequencing the whole complex and then bioinformatically deconvoluting the signal from the two cells (17, 18). This approach identifies which cell types are forming the complex but provides limited resolution of their transcriptional programs. While it is possible to manually dissect the doublets and then perform single-cell RNA sequencing, the process is long and fastidious and of limited throughput (18). Here, we designed an experimental workflow where cells forming complexes were isolated and physically separated from each other using fluorescence-activated cell sorting (FACS), followed by single-cell sequencing (**Figure 1A**).

**Figure 1:**
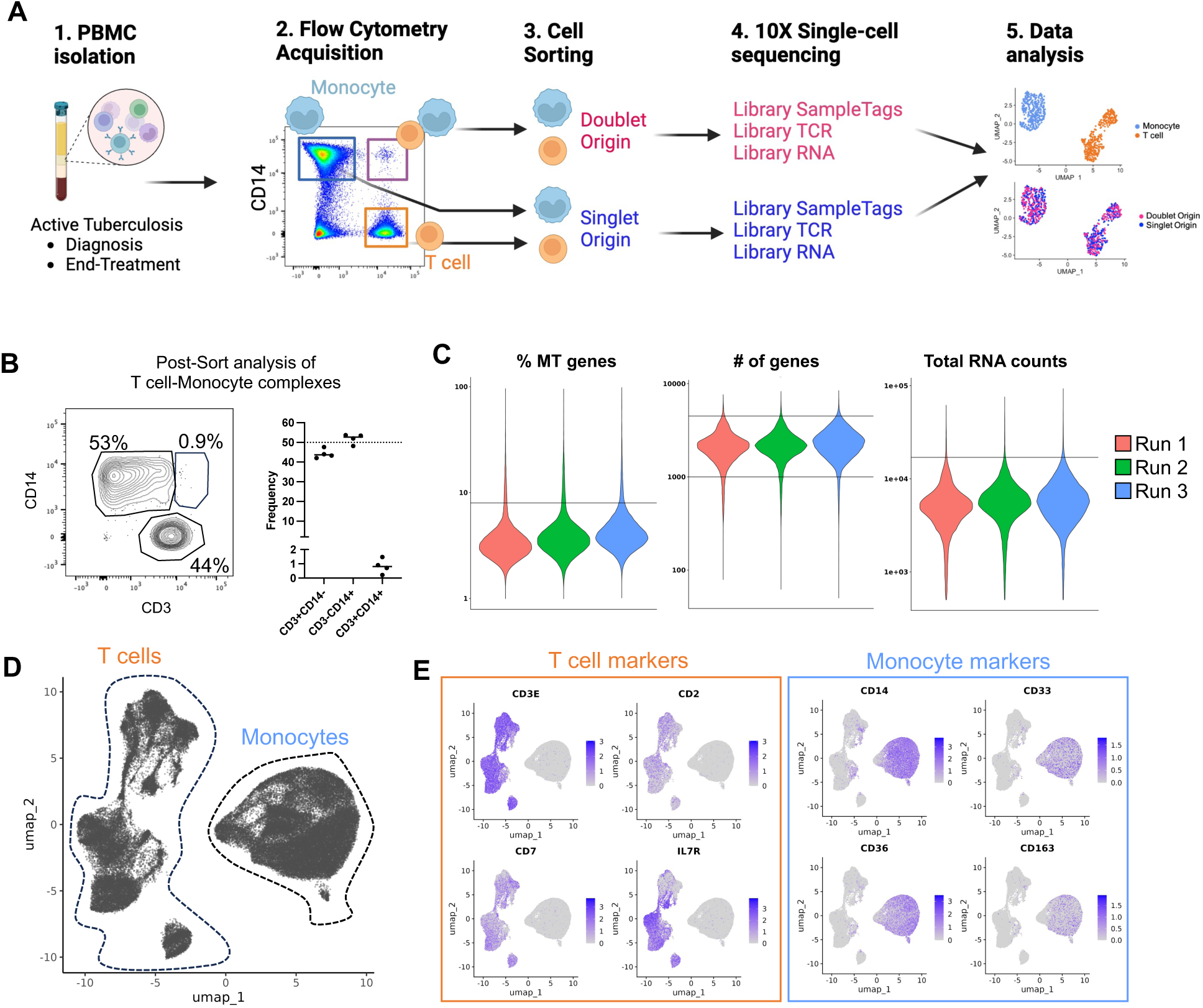
Determination of the single-cell transcriptome of thousands of T cells and monocytes forming complexes in ATB. A) Methodological workflow used to obtain the single-cell transcriptome and TCR repertoire of T cells and monocytes forming complexes (pink red, doublet origin) or singlet T cells and monocytes (blue, singlet origin) from cryopreserved human PBMC samples from ATB patients with samples collated at diagnosis and/or end-of-treatment, created with Biorender. B) Re-acquisition by flow cytometry of sorted T cell-Monocyte complexes in PBMC samples from four ATB patients at diagnosis. C) Percentage of mitochondrial genes, number of genes, and total RNA counts per cell across all three experimental runs. Black lines represent the thresholds used to remove suspected doublets and low-quality cells. D) Uniform manifold approximation and projection (UMAP) representation of all cells based on single-cell RNA reads. E) UMAP feature plot showing the expression level of canonical T cell and monocyte markers. Data were derived from 68,142 total cells, from 18 PBMC samples (Figure S1C).

T cell-monocyte complexes from cryopreserved peripheral blood mononuclear cells (PBMC) were sorted in bulk by flow cytometry as previously described (6) with the addition of CD19 exclusion (i.e., live CD3+CD14+CD19-events within the singlet gate, see **Figure S1A** for the full gating strategy). Cell sorting disrupts the physical connection between cells forming complexes, with the vast majority of cells being singlet CD3+ or singlet CD14+ cells post-sort (16). Here, we confirmed this observation by re-analyzing CD3+CD14+ sorted events from four PBMC samples and found an approximate 50/50 mix of CD3+CD14-singlet T cells and CD3-CD14+ singlet monocytes, with less than 2% dual positive CD3+CD14+ cells (**Figure 1B**). This post-sort single cell suspension was then used for droplet single-cell sequencing using the 10X genomics platform. In parallel, bulk-sorted singlet T cells and singlet monocytes mixed at a 1:1 ratio were run through the same droplet single-cell sequencing workflow but processed in separate libraries from the cells originating from complexes. Sequenced libraries were then integrated into one Seurat object. Thus, the resulting combined UMAP analysis showed two distinct cell types: T cells and monocytes, and depending on their library of origin, cells could also be additionally labeled as doublet origin (DO) or singlet origin (SO).

Using this experimental workflow, we processed PBMC samples from eight ATB patients who provided samples at diagnosis and after anti-TB therapy at six months post-diagnosis (i.e., end-of-treatment sample), and an additional two ATB patients with either a diagnosis sample or an end-of-treatment sample (**Table S1**). The majority of cells did not show a signature indicative of low quality (i.e., high frequency of mitochondrial genes, or low number of genes/RNA counts detected per cell), or intra-individual doublets (i.e., high number of genes or RNA counts per cell) (**Figure 1C**). After filtering out low-quality cells and doublets, we obtained a total of 68,142 single cells, of which 9,915 were DO (**Figure 1C**). After integration, we observed no batch effect between the three experimental runs (**Figure S1B**). As expected, UMAP clustering (**Figure 1D**) and expression of T cell and monocyte canonical markers (**Figure 1E**) identified two main groups of cells, corresponding to T cells (left side) and monocytes (right side). The UMAP clustering results were used to annotate each cell as either T cell or monocyte, and the total T cell and monocyte cell numbers recovered per sample are shown in **Figure S1C**. In conclusion, our novel experimental design defined the single-cell transcriptome of thousands of T cells and monocytes that were either singlets (i.e., SO) or forming complexes (i.e., DO).

### DO T cells are associated with a specific gene expression signature

A differential expression analysis using sample identifiers as a covariate (see methods) identified 193 genes upregulated in DO T cells and 72 genes upregulated in SO T cells (**Figure 2A**, **Table S2**). Upregulated genes in SO T cells were predominantly genes associated with translation (i.e., ribosomal genes) and genes associated with naïve T cells (CCR7, LEF1, TCF7) (**Figure 2A, 2B, Table S3A**). For genes upregulated in DO T cells, the top 10 GO terms were associated with inflammatory response (defense response, NFkB signaling, inflammatory response), T cell activation, MHC-II antigen presentation, and cell adhesion (**Figure 2C, Table S3B**). In addition, the top 50 genes upregulated in DO T cells encompassed several cytotoxicity genes (CST7, GZMB, NKG7, PRF1) (**Figure 2D**). Thus, DO T cells hold a unique gene expression signature characterized by several immune synaptic features such as MHC-II complex, NFkB signaling, cell adhesion, as well as effector T cell features (i.e., inflammation, activation, cytotoxicity).

**Figure 2:**
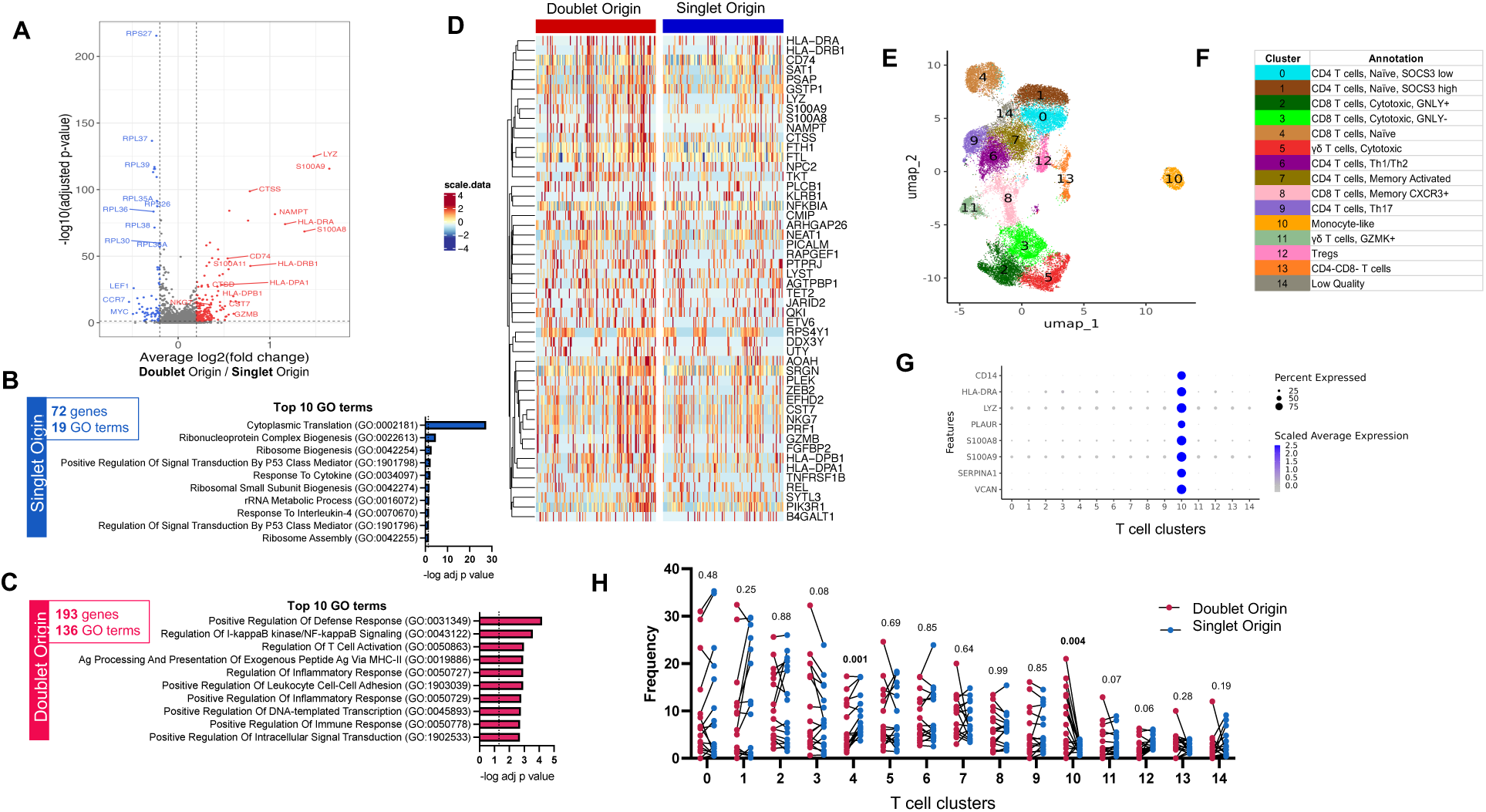
DO T cells are associated with a specific gene expression signature. A) Volcano plot of differentially expressed genes (DEGs) comparing doublet origin (DO) versus singlet origin (SO) T cells. Red dots represent DEGs upregulated in DO T cells (adjusted p-value < 0.05, average log2 fold change > 0.2), and blue dots represent DEGs upregulated in SO T cells (adjusted p-value < 0.05, average log2 fold change < −0.2). P-values were adjusted based on Bonferroni correction. B) Top 10 GO terms significantly associated with the 72 genes upregulated in SO T cells. C) Top 10 GO terms significantly associated with the 193 genes upregulated in DO T cells. D) Heatmap representation of the top 50 DEGs upregulated in DO T cells. Each column represents one DO or SO T cell. Color scale denotes RNA expression level after scaling. For visualization, SO T cells were randomly downsampled to have the same sample size as DO T cells. E) UMAP representation and F) manual cluster annotation of DO and SO T cells. G) Monocyte gene expression across all T cell clusters. H) T cell cluster composition differences between DO (red dots) and SO (blue dots) T cells paired by sample, using non-parametric paired Wilcoxon tests. Data were derived from 3,285 DO and 26,957 SO T cells, from 17 PBMC samples (Figure S1C).

A UMAP clustering analysis on all T cells (**Figure 2E**) was manually annotated (**Figure 2F**) based on top expressed genes per cluster (**Figure S2A**, **Table S4**). Clusters for naïve and memory CD4 and CD8 αβ T cells, cytotoxic αβ T cells, γδ T cells, Tregs, and double negative (DN) T cells were identified (**Figure 2F**). In addition, there was one outlier cluster with a strong monocyte signature (cluster 10, **Figure 2E** and **2G**). Cells from this cluster were assigned as T cells in the global UMAP analysis containing both T cells and monocytes (**Figure 1D**), indicating their transcriptomic profile is more similar to T cells than monocytes. However, unlike the majority of T cells, they also have expression of monocyte genes.

Next, we compared the cell cluster composition of DO versus SO T cells in each sample. Cluster 10, the outlier cluster with a monocyte signature, showed a striking enrichment for DO T cells (*p* = 0.004) (**Figure 2H**). Cluster 3, annotated as GNLY-negative cytotoxic CD8 T cells, was the second most enriched cluster in DO T cells, although not significant (*p* = 0.08). This matches our DE analysis result, where cytotoxic genes were found upregulated in DO T cells, as well as several defense response genes typically associated with monocytes: LYZ, NAMPT, S100A8, and S100A9 (**Figure 2D**). Only one cluster was at significantly increased prevalence in SO cells, cluster 4, representing naïve CD8 T cells (**Figure 2H**), also matching the DE analysis results. Together, our results indicate that effector T cells are preferentially forming immune synapses with monocytes in blood, possibly through TCR/MHC-II mediated interactions.

### DO monocytes are associated with a specific gene expression signature

The same analytical workflow identified fewer transcriptomic differences between DO and SO monocytes, namely 21 versus 19 genes upregulated in DO and SO categories, respectively (**Figure 3A**, **Table S5**). In genes upregulated in SO monocytes, only two GO terms were significantly enriched, with p-values close to the significance threshold (*p* = 0.045, **Figure 3B** and **Table S6A**). In contrast, in genes upregulated in DO monocytes several GO terms were enriched at high significance (*p* < 0.0001), related to MHC-II complex and cell adhesion (**Figure 3C, 3D** and **Table S6B**). Importantly, the immune synaptic myeloid cell adhesion molecule ICAM1 was upregulated in DO monocytes (**Figure 3A** and **3D**). In the same samples, HLA-DR protein expression was significantly higher in T cell-monocyte complexes compared to singlet T cells and monocytes (p < 0.0001, **Figure 3E**), corroborating our finding that several MHC-II genes were upregulated in both DO T cells (**Figure 2A**) and monocytes (**Figure 3A**).

**Figure 3:**
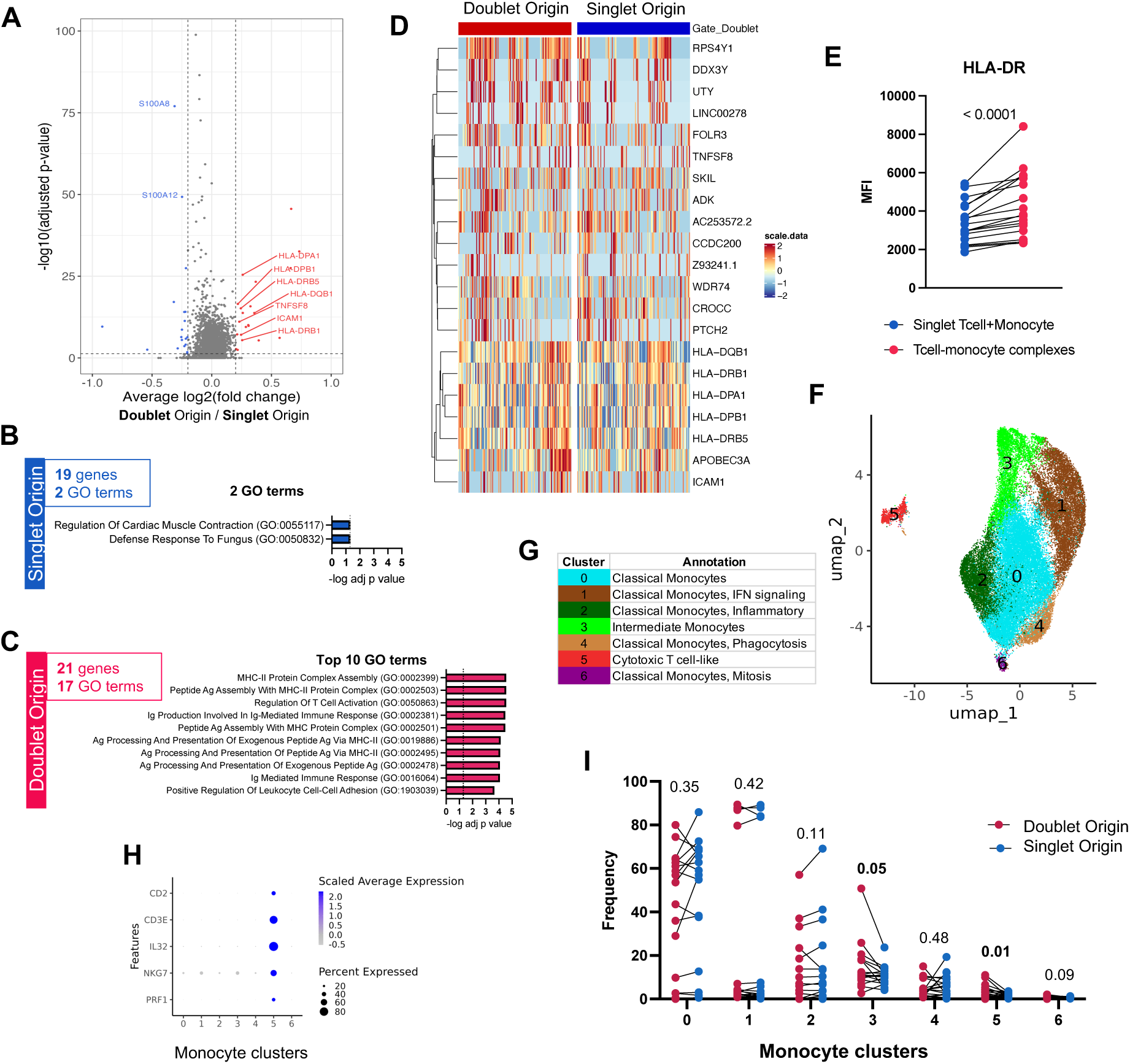
DO monocytes are associated with a specific gene expression signature. A) Volcano plot of DEGs comparing monocytes of doublet versus SO. Red dots represent DEGs upregulated in doublet origin (DO) monocytes (adjusted p-value < 0.05, average log2 fold change > 0.2), blue dots represent DEGs upregulated in singlet origin (SO) monocytes (adjusted p-value < 0.05, average log2 fold change < −0.2). P-values were adjusted based on Bonferroni correction. B) GO terms significantly associated with the 19 genes upregulated in SO monocytes. C) Top 10 GO terms significantly associated with the 21 genes upregulated in DO monocytes. D) Heatmap representation of 21 DEGs upregulated in DO monocytes. Each column represents one DO or SO monocyte. Color scale denotes RNA expression level after scaling. For visualization, SO monocytes were randomly downsampled to have the same sample size as DO monocytes. E) Median fluorescence intensity (MFI) of HLA-DR protein expression on the surface of T cell-monocyte complexes (T-M), or the sum of singlet T cells and singlet monocytes (T+M). All three populations were defined by flow cytometry as shown in Figure S1A. F) UMAP representation and G) manual cluster annotation of DO and SO monocytes. H) Cytotoxic T cell gene expression across all monocyte clusters. I) Monocyte cluster composition differences between DO (red dots) and SO (blue dots) monocytes paired by sample, using non-parametric paired Wilcoxon tests. Data were derived from 5,530 DO and 31,270 SO monocytes, from 17 PBMC samples (Figure S1C).

A cell subset composition analysis of monocytes was performed as described for T cells (**Figure 3F**), with manual annotation of each cluster (**Figure 3G**) based on their top expressed genes (**Figure S2B**, **Table S7**). We identified several clusters of classical monocytes associated with distinct cellular processes (i.e., interferon signaling, inflammation, phagocytosis, mitosis), and one cluster of intermediate monocytes with FCGR3A (CD16) and high MHC-II expression (**Figure 3G**). Strikingly, and mirroring our T cell analysis, there was one outlier cluster with high expression of cytotoxic T cell genes (cluster 5, **Figure 3F** and **Figure 3H**). Cells from this cluster were assigned as monocytes in the global UMAP analysis containing both T cells and monocytes (**Figure 1D**), indicating their transcriptomic profile is more similar to monocytes than T cells. However, unlike the majority of monocytes, they also have expression of T cell genes.

When comparing the cell cluster composition between DO and SO monocytes, no cluster had a higher prevalence in SO cells (**Figure 3I**). In contrast, two clusters had a significantly higher prevalence in DO cells: cluster 3, corresponding to intermediate monocytes (*p* = 0.05), and the cytotoxic T cell-like cluster 5 (*p* = 0.01) (**Figure 3I**). Together, our results demonstrate that DO monocytes hold a unique transcriptomic signature associated with immune synaptic components (i.e., MHC-II complex, ICAM1), and are enriched for intermediate and cytotoxic T cell-like monocyte subsets.

### Outlier DO T cells and monocytes are separate entities

To confirm that the monocyte-like cluster 10 in the T cell UMAP analysis (**Figure 2E**) and the cytotoxic T cell-like cluster 5 in the monocyte UMAP analysis (**Figure 3F**) indeed represented separate entities and not one single T cell-monocyte dual expressing population, we ran a UMAP clustering analysis combining the two outlier clusters. We found three clusters, clearly separating T cells (clusters 0 and 1) from monocytes (cluster 2) (**Figure S2C** and **S2D**). The two T cell clusters represented cytotoxic T cells (cluster 0) and naïve T cells (cluster 1), and the monocyte cluster (cluster 2) was associated with inflammatory monocytes (**Figure S2E**). Monocyte-like DO T cells were enriched for the cytotoxic cluster 0, whereas a 50/50 mix was found for monocyte-like SO T cells (**Figure S2F**). Thus, the T cell-monocyte dual positive clusters within the T cell and the monocyte UMAP analyses are separate entities, representing either cytotoxic/naive T cells co-expressing monocyte genes, or inflammatory monocytes co-expressing T cell genes, respectively. In addition, monocyte-like DO T cells were enriched for cytotoxic over naïve phenotypes.

### Increased immune activation in DO T cells and monocytes in ATB at diagnosis

Next, we investigated differences between diagnosis and end-of-treatment complexes. DO T cell and DO monocyte gene signatures were similarly expressed in DO cells between diagnosis and end-of-treatment (**Figure S3A** and **S3B**). In terms of cell subsets, no differences in cell cluster composition were found between paired diagnosis and end-of-treatment samples in DO (**Figure S3C**) or SO T cells (**Figure S3D**). In monocytes, both DO and SO cells showed enrichment for interferon-signaling classical monocytes (cluster 1) at diagnosis and enrichment for inflammatory classical monocytes (cluster 2) at end-of-treatment (**Figure S3E** and **S3F**). In addition, DO monocytes showed enrichment for intermediate monocytes (cluster 3) at end-of-treatment (**Figure S3E**). Thus, the transcriptomic signature and outlier cluster enrichment in DO T cells and monocytes described in the sections above were unaffected by disease resolution, indicating that they represent core features of T cell-monocyte complexes.

To specifically investigate disease-related transcriptomic differences associated with T cells and monocytes forming complexes, we performed a paired differential expression analysis between diagnosis and end-of-treatment DO or SO cells. Hundreds of genes were upregulated at diagnosis compared to end-of-treatment in DO or SO cells, with a significant overlap (324 genes in DO T cells, including 105 shared (32%) with SO T cells, **Figure 4A**; 743 genes in DO monocytes, including 468 shared (63%) with SO monocytes, **Figure 4F**).

**Figure 4:**
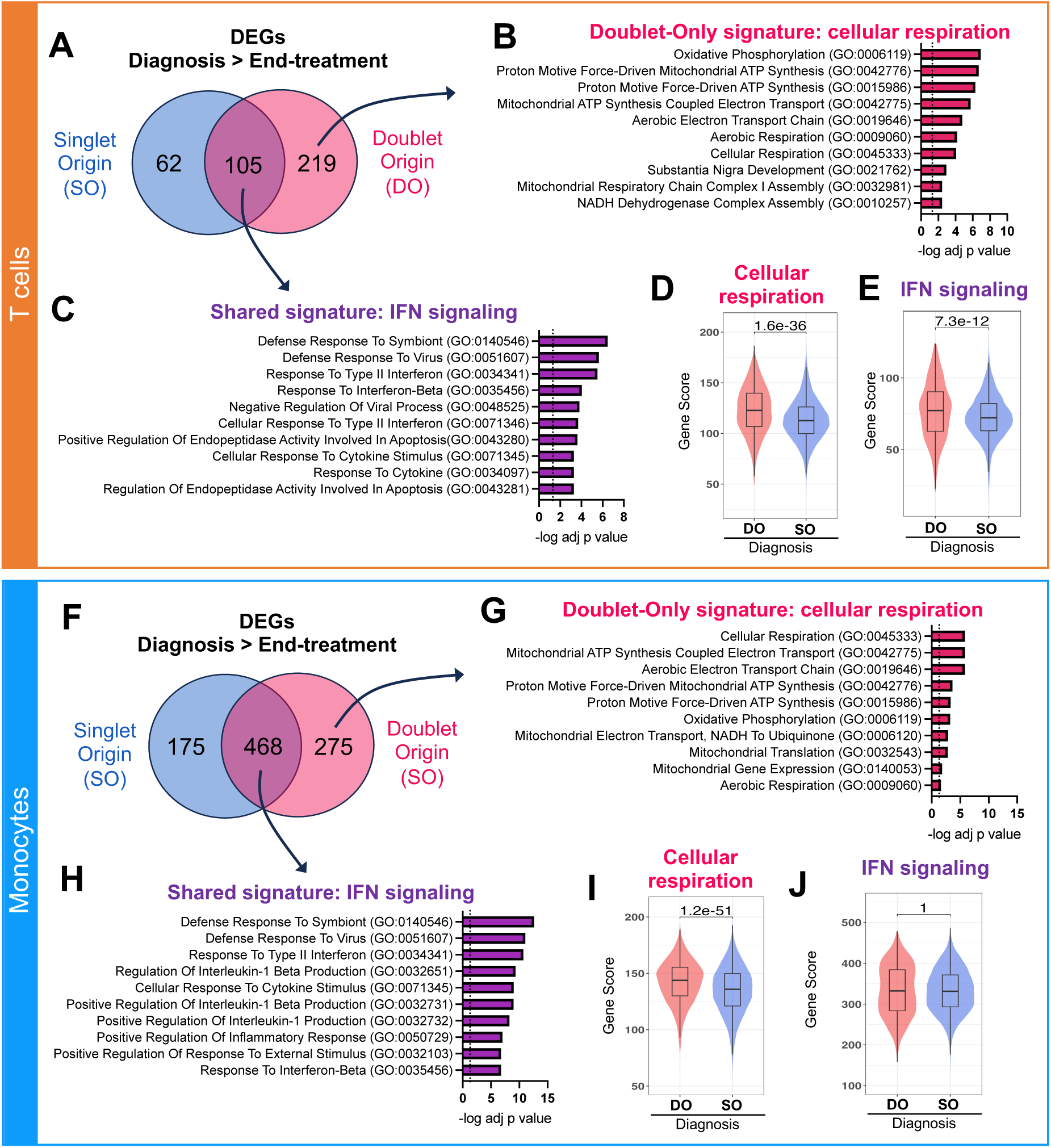
Increased immune activation in DO T cells and monocytes in ATB at diagnosis. A) Overlap between upregulated DEGs (adjusted p-value with Bonferroni correction <0.05, average log2 fold change > 0.2) at diagnosis versus end-of-treatment in singlet origin (SO, blue) and doublet origin (DO, pink red) origin T cells. B) Top 10 GO terms significantly associated with the 219 DEGs upregulated at diagnosis in DO but not SO T cells (i.e., doublet-only signature), indicating a strong association with cellular respiration. C) Top 10 GO terms significantly associated with the 105 genes upregulated at diagnosis in both SO and DO T cells (i.e., shared signature), indicating a strong association with IFN signaling. Distribution of the D) cellular respiration gene signature score and E) IFN signaling gene signature score in DO versus SO T cells at diagnosis. The cellular respiration gene signature score represents the sum expression of the 219 DEGs upregulated at diagnosis versus end-of-treatment in DO but not SO T cells, as defined in A. The IFN signaling gene signature score represents the sum expression of the 105 DEGs upregulated at diagnosis versus end-of-treatment in both SO and DO T cells, as defined in A. F) Overlap between upregulated DEGs (adjusted p-value with Bonferroni correction <0.05, average log2 fold change > 0.2) at diagnosis versus end-of-treatment in singlet origin (SO, blue) and doublet origin (DO, pink red) origin monocytes. G) Top 10 GO terms significantly associated with the 275 DEGs upregulated at diagnosis in DO but not SO monocytes (i.e., doublet-only signature), indicating a strong association with cellular respiration. C) Top 10 GO terms significantly associated with the 468 genes upregulated at diagnosis in both SO and DO monocytes (i.e., shared signature), indicating a strong association with IFN signaling. Distribution of the D) cellular respiration gene signature score and E) IFN signaling gene signature score in DO versus SO monocytes at diagnosis. The cellular respiration gene signature score represents the sum expression of the 275 DEGs upregulated at diagnosis versus end-of-treatment in DO but not SO monocytes, as defined in A. The IFN signaling gene signature score represents the sum expression of the 468 DEGs upregulated at diagnosis versus end-of-treatment in both SO and DO monocytes, as defined in A. For the boxplots in D-E) and I-J), the lower, median, and upper edges represent the 25th, 50th, and 75th percentile; the length of the upper and lower whiskers is 1.5 times the interquartile range. Non-parametric unpaired Mann-Whitney tests were used for comparison between DO and SO cells, and Bonferroni correction was performed to adjust the p-value. Data were derived from seven diagnosis/end-of-treatment PBMC sample pairs, one unpaired diagnosis sample, and one unpaired end-of-treatment sample (Figure S1C).

Genes upregulated at diagnosis in DO but not SO cells (i.e., doublet-only signature) were associated with cellular respiration in both T cells (**Figure 4B**) and monocytes (**Figure 4G**). At diagnosis, the cellular respiration signature was significantly upregulated in DO versus SO T cells (**Figure 4D**) and monocytes (**Figure 4I**). Genes upregulated at diagnosis compared to end-of-treatment in both DO and SO cells (i.e., shared signature) were associated with type 1 and type 2 interferon (IFN) signaling in both T cells (**Figure 4C**) and monocytes (**Figure 4H**). At diagnosis, the IFN signature was significantly upregulated in DO versus SO T cells (**Figure 4E**) but not monocytes (**Figure 4J**). Thus, in ATB disease, the transcriptomic signature of T cells and monocytes forming complexes at diagnosis indicated higher cellular respiration compared to their singlet counterparts. In addition, at diagnosis, DO T cells showed increased expression of IFN signaling genes compared to SO T cells.

### Active TCR signaling in DO T cells in ATB

We have previously shown that in steady-state, T cell-monocyte complexes showed LFA1/ICAM1 polarization but not CD3 polarization, suggesting that they were not mature immune synapses (6). To determine whether this was also the case during ATB disease, we examined CD3 polarization in T cell-monocyte complexes from PBMC collected at diagnosis in two ATB participants using confocal microscopy. Complexes were fixed before sorting to retain their integrity. In both patients, over 70% of T cell-monocyte complexes showed CD3 polarization at the point of contact (8 out of 11 complexes for patient A, and 7 out of 10 for patient B, **Figure 5A**). In contrast, less than 15% of T cell-monocyte complexes isolated from PBMC of a Mtb-negative participant using the same protocol displayed such a pattern (3 out of 23, **Figure 5A**). In an additional four ATB participants with PBMC samples at diagnosis, using flow cytometry, we found a marked higher protein expression of TCRαβ but not TCRγδ in T cell-monocyte complexes compared to singlet T cells and singlet monocytes combined (**Figure 5B**). Thus, during ATB disease, T cells forming complexes present several features indicative of active TCR signaling, namely CD3 polarization at the point of contact with monocytes and higher TCRαβ expression.

**Figure 5:**
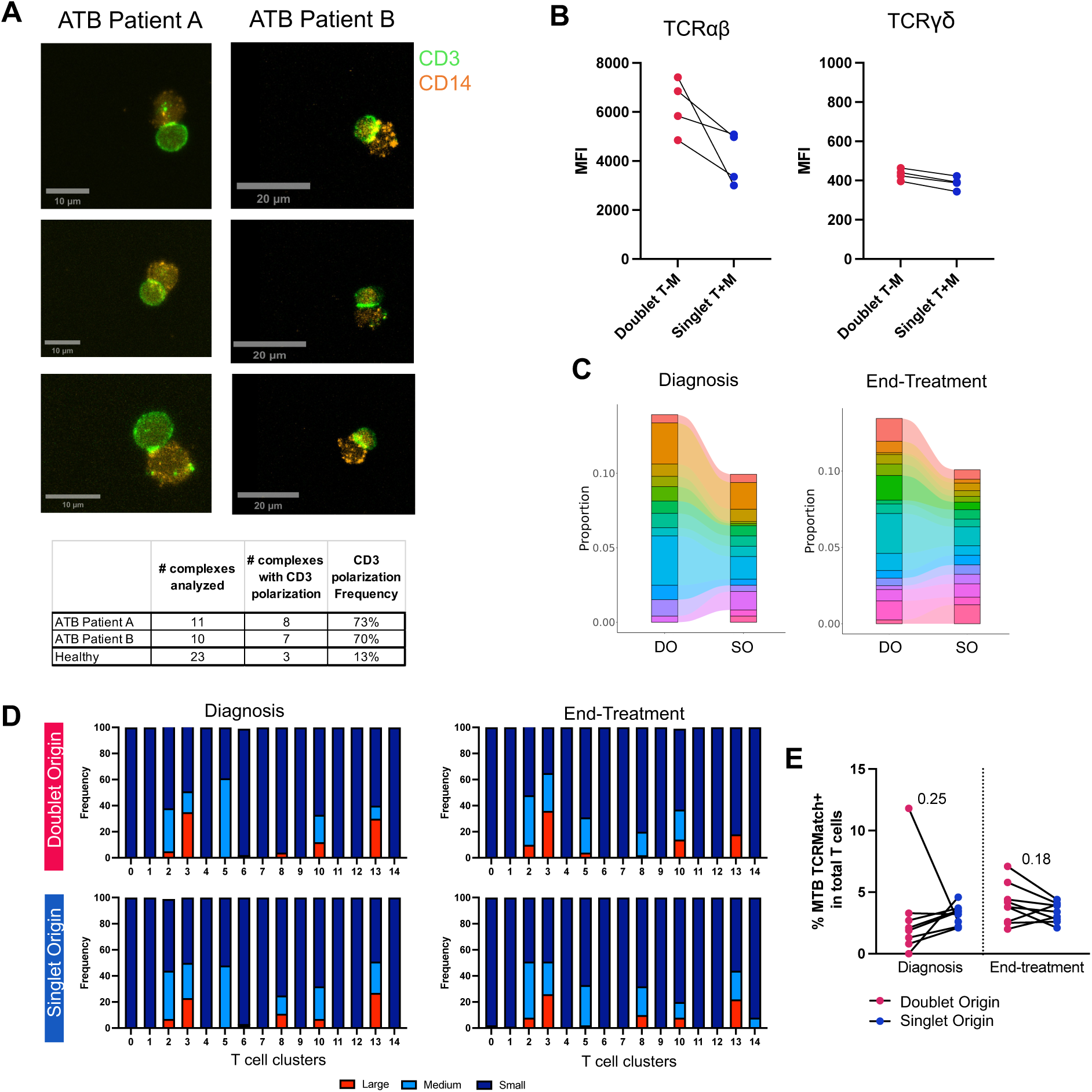
Active TCR signaling and higher clonal expansion in DO T cells in ATB. A) Representative images of CD3 and CD14 expression on fixed T cell-monocyte complexes isolated by cell sorting from PBMC samples collected at diagnosis from two patients with ATB. The bottom table represents the number of complexes analyzed, and the number of complexes with CD3 polarization per sample. B) Median fluorescence intensity (MFI) of TCRαβ and TCRγδ protein expression on the surface of T cell-monocyte complexes, or the sum of singlet T cells and singlet monocytes, in PBMC samples from four ATB patients at diagnosis. All three populations were defined by flow cytometry as shown in Figure S1A. C) Frequency of the top 10 TCR clonotypes by abundance in DO and SO T cells at diagnosis, and end-of-treatment (since there was a tie when ranking TCR clonotypes by proportion of total T cells, the top 10 correspond to 14 and 16 individual TCR clonotypes for diagnosis and end-of-treatment samples, respectively). D) Frequency of small, medium, and large TCR clonotypes in doublet and SO T cells at diagnosis and end-of-treatment, per individual T cell cluster (as defined in Figure 2E). Small denotes clonotypes with cell count <= 5, medium denotes clonotypes with 5 < cell count <= 20, and large denotes clonotypes with cell count > 20. E) Frequency of T cells expressing a Mtb-specific TCRβ CDR3 sequence (defined with TCRMatch (25)) in DO versus SO T cells, at diagnosis and end-of-treatment. For C-E, data were derived from 24,025 T cells with both TCRα and TCRβ chains detected, from seven diagnosis/end-of-treatment PBMC sample pairs, one unpaired diagnosis sample, and one unpaired end-of-treatment sample (Figure S1E).

### Higher clonal expansion in DO T cells

In parallel, we compared the TCRαβ repertoire of DO and SO T cells. TCRαβ were the TCR-coupled chains expressed by the majority of DO T cells at the protein (**Figure 5B**) and gene level (**Figure S1E**). In both diagnosis and end-of-treatment samples, the most abundant TCRαβ clonotypes were almost entirely shared between DO and SO T cells; but in DO T cells, they represented a higher fraction of total cells, indicating higher clonal expansion (**Figure 5C**).

In both DO and SO T cells, large clones were restricted to five clusters: the cytotoxic T cell clusters 2 and 3, the CXCR3+ memory CD8 T cell cluster 8, the monocyte-like T cell cluster 10, and the DN T cell cluster 13 (**Figure 5D**). In cluster 3 and cluster 10 (the two clusters at increased prevalence in DO T cells), the frequency of large clones was higher in DO compared to SO T cells (**Figure 5D**). The proportion of large clones within each cluster remained largely unchanged between diagnosis and end-of-treatment samples for both DO and SO T cells, indicating that the higher clonal expansion in DO T cells was independent of the presence of active infection (**Figure 5D**).

Finally, we explored the antigen specificity of the TCR sequences retrieved in DO and SO T cells. TCRMatch is a publicly available online tool that predicts TCR antigen-specificity based on previously identified TCRs with known epitope specificity curated in the Immune Epitope Database (IEDB) (25). Using TCRMatch, we found positive matches to Mtb in DO T cells in 16 of the 17 samples analyzed, and at a similar frequency to SO T cells in both diagnosis and end-of-treatment samples (**Figure 5E**). Together, our results indicate that the TCR repertoire largely overlapped between DO and SO T cells, with the presence of antigen-specific T cells in both groups. However DO T cells were associated with a higher clonal expansion, a feature of effector T cells.

### Circulating T cell-monocyte complexes in dengue hold similar transcriptomic signatures

Finally, we applied the same strategy to separate cells forming complexes and performed single-cell sequencing (**Figure 1A**) in another infection system where we previously reported the presence of T cell-monocyte complexes: dengue (6). We studied a set of 15 PBMC samples of patients with dengue, collected in the acute (four to five days since symptom onset) and/or convalescent (14 to 21 days since symptom onset) phase of infection. After QC filtering, we recovered a total of 2,434 DO cells and 3,335 SO cells, including six samples with paired DO and SO cells (**Figure S4A**). Similar to the ATB dataset, T cells and monocyte**s were** clearly separated in the UMAP analysis (Figure S4B), based on their top 10 expressed genes (**Figure S4C**).

We found 89 genes upregulated in DO T cells, associated with 62 GO terms (**Figure S4D, Table S8**). The top 10 enriched GO terms included cytokine signaling, cell adhesion, and viral and innate immunity (**Figure 6A**). Within the 89 genes significantly upregulated in DO T cells, 20 overlapped with the T cell doublet signature found in ATB (i.e., T193 signature, red dots in Figure 2A), including the cytotoxic gene CTSS and the NFkB-related gene NFKBIA (**Figure 6B**, statistical significance of overlap *p* = 8e-25). Several activation markers were additionally present in DO T cells in dengue, such as CD69, STAT3, STAT4, TNFAIP3, and the cell-adhesion-related chemokine receptor CXCR4 (**Figure S4D**, **Table S8**). The T193 signature was also significantly upregulated in DO compared to SO T cells in the dengue acute but not convalescent-phase samples (**Figure 6C**). In DO monocytes, 148 genes were upregulated and associated with 143 GO terms (**Figure S4E, Table S9**). The top 10 enriched GO terms included several associated with MHC-II complex and one with cell adhesion (**Figure 6D**). Within the 148 genes significantly upregulated in DO monocytes, seven genes overlapped with the monocyte doublet signature found in ATB (i.e., M21 signature, red dots in Figure 3A), including MHC-II complex genes HLA-DPA1, HLA-DPB1 and HLA-DQB1 (**Figure 6E**, statistical significance of overlap *p* = 1e-11). Several other MHC-II genes were also upregulated in dengue DO monocytes (CD74, HLA-DRA, HLA-DMA, HLA-DQA1). The M21 signature was also significantly upregulated in DO compared to SO monocytes in the dengue dataset in both acute and convalescent PBMC samples (**Figure 6F**). Thus, we identified similarities in the transcriptome of T cells and monocytes forming complexes between ATB and dengue, in particular upregulation of genes associated with MHC-II complex, cell adhesion, and T cell activation.

**Figure 6:**
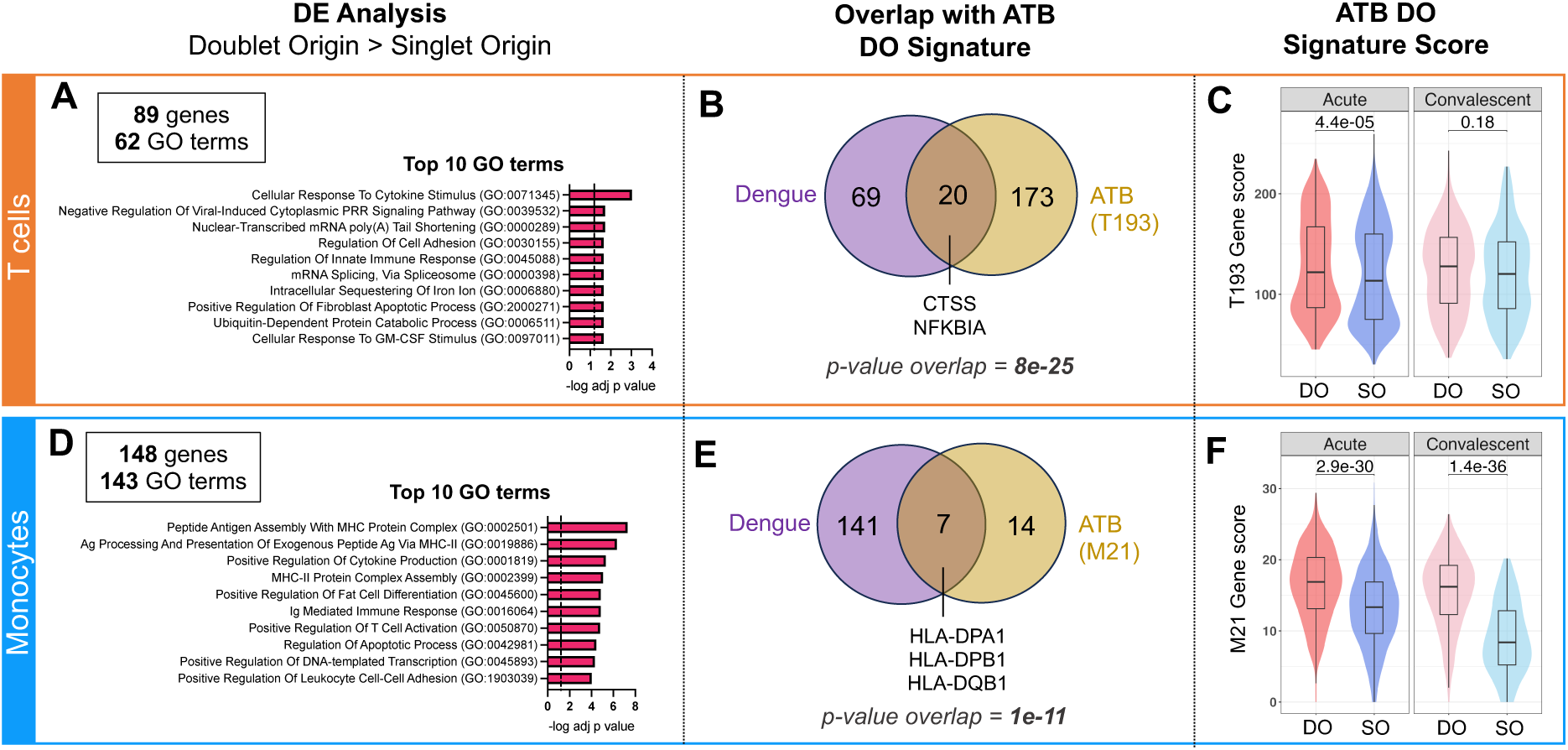
Circulating T cell-monocyte complexes in dengue hold similar transcriptomic signatures to ATB. The single-cell transcriptome of T cells and monocytes forming complexes (DO) or singlet T cells and singlet monocytes (SO) from 15 cryopreserved human PBMC samples from patients with dengue, with blood collected at either the acute (four to five days since symptom onset) or convalescent phase (14 to 21 days since symptom onset) were obtained by following the same workflow as for ATB (Figure 1A). Differential expression analysis between DO and SO cells was performed on T cells (*top panel*) and Monocytes separately (*bottom panel*). A) Top 10 GO terms significantly associated with the 89 genes significantly upregulated (adjusted p-value with Bonferroni correction <0.05, average log2 fold change > 0.2) in DO versus SO T cells. B) Overlap between genes significantly upregulated in DO versus SO T cells in dengue (purple) and ATB (yellow). The ATB signature of DO T cells (also referred to as T193) represents the 193 genes significantly upregulated in DO versus SO T cells in the ATB dataset, as defined in Figure 2A. C) Distribution of the T193 gene signature score in DO versus SO T cells in the dengue dataset, separating acute and convalescent samples. D) Top 10 GO terms significantly associated with the 148 genes significantly upregulated (adjusted p-value with Bonferroni correction <0.05, average log2 fold change > 0.2) in DO versus SO monocytes. E) Overlap between genes significantly upregulated in DO versus SO monocytes in dengue (purple) and ATB (yellow). The ATB signature of DO monocytes (also referred to as M21) represents the 21 DEGs upregulated in doublet versus SO T cells in the ATB dataset, as defined in Figure 3A. F) Distribution of the M21 gene signature score in DO versus SO monocytes in the dengue dataset, separating acute and convalescent samples. Non-parametric unpaired Mann-Whitney U tests were used for comparison between DO and SO cells, and Bonferroni correction was performed to adjust the p-value. Data were derived from 15 PBMC samples (Figure S4A).

## Discussion

This study describes the first single-cell transcriptomic analysis of immune cells forming complexes in human blood. We developed a novel high-throughput experimental workflow that allows for the isolation and physical separation of cells forming complexes in an automated fashion using FACS, followed by single-cell sequencing. This method can be applied to study thousands of cells forming complexes in a single sample and bypass the need for bioinformatic deconvolution required when analyzing complexes as a whole. This workflow can be used for the study of any type of immune cell complexes as long as each cell component expresses one distinct protein on the cell surface that can be detected by flow cytometry. It can also be combined with any single-cell sequencing technique that uses a single cell suspension as the starting material.

Applying this workflow to a cohort of PBMC samples from ATB patients collected at diagnosis and after anti-TB therapy at six months post-diagnosis, we found that the transcriptomic signature of T cells and monocytes forming complexes was associated with many TCR/MHC immune synaptic components, including MHC-II complex, cell adhesion, and NFkB signaling. In addition, at diagnosis, we found that the transcriptome of T cells and monocytes forming complexes indicated higher immune activation and metabolic activity compared to post-treatment, especially for T cells; and gene, TCR, and imaging features indicative of active TCR signaling. Thus, our method allowed the discovery of unique immune signatures in T cells and monocytes forming complexes, indicating that they engage in active TCR/MHC immune synapses during infection.

In addition, the transcriptomic signature of T cells and monocytes forming complexes was associated with cytotoxic T cells and MHC-II^high^ intermediate monocytes. We have recently shown that amongst circulating monocyte subsets, CD14+CD16+ intermediate monocytes showed the highest transcriptomic changes in ATB at diagnosis, with upregulation of genes associated with MHC-II complex and inflammation (26). In T cells, where Th1 and Th1* phenotypes are typically associated with Mtb protective immunity (27–29), there is growing evidence that cytotoxic T cells also play a role in TB infection (30–32). Thus, by providing a snapshot of immune cells actively interacting at the time of blood draw, circulating T cell-monocyte complexes may uncover novel T cell and monocyte subsets that play an active role in the immune response to infection.

Unexpectedly, we found that DO cells were markedly enriched in two outlier clusters: one composed of T cells expressing monocyte genes and another of monocytes expressing T cell genes. These clusters exhibited the highest enrichment in DO compared to SO cells. We provided experimental and computational evidence to confirm that these cells represented separate entities (i.e., either T cells or monocytes), and were singlets, not intact cell-cell complexes. In addition, monocyte-like T cells and T cell-like monocytes displayed phenotypic similarities, with elevated expression of MHC-II and cytotoxic genes. Thus, an intriguing implication from these findings is that RNA exchange occurs between T cells and monocytes forming complexes. This process has been observed during cell-cell interactions through exosomes (33), and also direct contact using membrane protrusions such as nanotubes (34). In addition, the exchange of protein and RNA material has been demonstrated at the immune synapse, through microvesicles (35–37) and exosomes (38). Additional research will be needed to understand the precise mechanism of RNA transfer within circulating T cell-monocyte complexes, possibly using high-resolution microscopy. Regardless of the mechanism by which RNA is exchanged between cells forming complexes, its significance is large. First, the retained RNA footprint could be used to monitor recent physical interactions between immune cells. This concept has been already proposed at the protein level to monitor interactions in tissues between T cells and B cells (39) and between CD8 T cells and myeloid cells (40). More recently, a neighboring cell analysis study found that cells in tissues share similar transcriptomic signatures to their neighboring cells, indicating the occurrence of RNA transfer following interactions (41). Here we provide seminal evidence that it may occur as well between interacting cells in blood. The second implication relates to doublet detection algorithms, which typically identify heterotypic doublets (i.e., a complex of two cells from a distinct lineage) based on the co-expression of gene programs that are specific to one single cell type (42). It is unlikely that these algorithms will be able to distinguish between heterotypic doublets and singlet cells that have recently interacted with another immune cell type and received some of their RNA. Indeed, several single-cell transcriptomic studies have shown the presence of singlet cells with dual lineage expression signatures that resemble doublets, even after applying doublet detection algorithms (13–15, 43).

Finally, we found that the transcriptome of T cell-monocyte complexes in individuals with dengue significantly overlap with those from ATB, including genes associated with T cell activation, cell adhesion, and MHC-II. Thus, the presence of TCR/MHC-II immune synapses between T cells and monocytes in blood may be a common feature during infection. Since circulating T cell-monocyte complexes have also been described in many other immune perturbation models, including vaccination (6), cancer (10, 11), and chronic inflammation (12), it would be extremely valuable to check whether the same signatures hold, or if other mechanisms are at play in distinct immune perturbation contexts.

Our study has several limitations. First, our method does incur a loss of pairing between cells forming complexes. It is possible to infer which cell types were likely interacting based on their shared RNA signatures (in our case, cytotoxic T cells and MHC-II^high^ monocytes), but this ability will be impaired if more than one cell subset of T cells or monocytes are interacting with each other. Thus, this method may be even more informative when used in conjunction with other high-throughput whole complex single-cell techniques such as PIC-Seq (17) to reconnect interacting cell subsets. Second, as for all other methods studying cell-cell complexes, our method is still confounded by the fact that not all CD3+CD14+ events detected in human blood are biological T cell-monocyte complexes. Many are expected to be technical artifacts, which coincidentally were too close to each other to be detected as a doublet by the cell sorter. Our dataset reflects this caveat, particularly in T cells, where the majority of transcriptomic differences between DO and SO cells were found in only a handful of clusters. These clusters likely represent T cells forming biological synapses, with the remaining clusters representing “noise” from coincidental interactions. Distinguishing between synaptic versus coincidental doublets, for instance by using imaging features from recently developed high-throughput imaging sorting technologies (44), should further increase the resolution of our method.

In conclusion, we developed a novel method to study the single-cell transcriptome of T cells and monocytes forming complexes from blood samples, that can be easily adapted for the study of any cell-cell interactions. Applying this method to ATB and dengue disease cohorts, we provided several compelling pieces of evidence that T cell-monocyte complexes in human blood represent active TCR/MHC-II immune synaptic interactions, with the most activity at the clinical phase of infection, and are enriched for T cells and monocytes subsets expected to play important functions during infection. We also found that within complexes, T cells showed more changes over monocytes and that RNA is exchanged between interacting cells, two valuable novel insights that would have been missed if studying complexes as a whole. Thus, studying the single-cell transcriptome of T cells and monocyte forming complexes in blood is a valuable strategy to monitor immune synaptic interactions during infection.

## Methods

### Ethics statement

Human study participants were enrolled at the South African Tuberculosis Vaccine Initiative, University of Cape Town, Western Cape Province (South Africa) for ATB, and in the Pediatric Dengue Hospital-based Study (Nicaragua) for dengue. Ethical approval to carry out this work was maintained through the La Jolla Institute for Immunology Institutional Review Board (IRB), the Human Research Ethics Committee of the University of Cape Town, the University of Colombo Ethics Review Committee, the Nicaraguan Ministry of Health, and the UC Berkeley Center for the Protection of Human Subjects. All clinical investigations were conducted according to the principles expressed in the Declaration of Helsinki, and all participants (or guardians for participants <18 years old) provided written informed consent before participation in the study. In Nicaragua, children 6 years and older provided assent.

### Study Cohorts and Samples

ATB and dengue cohorts’ descriptions and demographics are available in **Table S1**. ATB was defined as 1) the presence of clinical symptoms and/or radiological/histological evidence of pulmonary TB, and 2) microbiological confirmation by *Mtb*-specific molecular testing on sputum. For ATB subjects, blood samples were obtained at diagnosis and the end of a six-month anti-TB therapy. Anti-TB therapy was a standard regimen for drug-susceptible *Mtb* consisting of an intensive phase of two months with isoniazid (INH), rifampin (RIF), pyrazinamide (PZA), and ethambutol (EMB) followed by a continuation phase of four months with INH and RIF (45). Dengue samples were collected in the Hospital Infantil Manuel de Jesús Rivera (HIMJR) in Managua, the capital city of Nicaragua. Blood samples were obtained at the acute (four to five days since symptom onset), and convalescent (14 to 21 days since symptom onset) phases of infection. Dengue fever (DF) was defined as acute febrile illness with two or more of the following: headache, retro-orbital pain, myalgia, leukopenia, arthralgia, rash, and hemorrhagic manifestations. DHF was defined as DF with hemorrhagic manifestations, thrombocytopenia, and signs of plasma leakage (46). Peripheral blood mononuclear cells (PBMC) were obtained by density gradient centrifugation (Ficoll-Hypaque, GE Healthcare) from whole-blood samples, according to the manufacturer’s instructions. Cells were resuspended at up to 10 million cells per milliliter in FBS (Gemini Bio-Products) containing 10% DMSO (Sigma) and cryopreserved in liquid nitrogen. Healthy controls had no past medical history of TB, nor exposure to *Mtb* or evidence of *Mtb* sensitization as confirmed by a negative IFNγ–release assay. All participants were confirmed negative for human immunodeficiency virus (HIV) infection.

### PBMC thawing

Cryopreserved PBMC were quickly thawed by incubating each cryovial at 37°C for 2 min, and cells were transferred into 9 ml of cold medium (RPMI 1640 with L-Glutamine and 25 mM Hepes (Omega Scientific), supplemented with 5% human AB serum (GemCell), 1% Penicillin Streptomycin (Gibco), 1% Glutamax (Gibco)), and 20 U/mL Benzonase Nuclease (Millipore). Cells were centrifuged and resuspended in medium to determine cell concentration and viability using Trypan blue and a hematocytometer. Cells were then kept at 4°C until used for flow cytometry or cell sorting.

### Non-imaging Flow Cytometry Acquisition and Cell Sorting

After PBMC thawing, up to 10×10^6^ cells were surface stained with fluorochrome-conjugated antibodies, as previously described (47). Cells were incubated with 10% FBS in 1X PBS for 10 minutes. Cells were then stained with 100 μl of PBS containing 0.1 μl fixable viability dye eFluor506 (eBioscience, corresponding to 1:1000 dilution of the stock, as per the manufacturer’s recommendation), 2 μl of FcR blocking reagent (Biolegend), and various combinations of anti-human CD19 PE-Cy7 (2 μl per test, clone HIB19, TONBO Biosciences), CD3 AF488 (2 μl per test, clone UCHT1, Biolegend), CD3 AF700 (3 μl per test, clone UCHT1, eBiosciences), CD14 APC (2 μl per test, clone 61D3, TONBO Biosciences), CD14 BV421 (1 μl per test, clone M5E2, Biolegend), HLA-DR PE (2 μl per test, clone L243, Biolegend), TCRαβ PEdazzle594 (2 μl per test, clone IP26, Biolegend) and TCRγδ BV421 (2 μl per test, clone 11F2, BD Biosciences) for 20 min at room temperature. For samples that were used for single-cell sequencing, TotalSeqTM-C oligonucleotide-conjugated antibodies (Biolegend) were also added at this step at 0.01mg/mL final concentration (one distinct antibody per sample). After two washes in PBS, cells were resuspended into 100 μl of MACS buffer (PBS containing 2mM EDTA (pH 8.0) and 0.5% BSA) and stored at 4°C protected from light for up to four hours until flow cytometry acquisition. Cell sorting was performed on a BD FACSAria Fusion cell sorter (Becton Dickinson). T cell-monocyte complexes, singlet T cells, and singlet monocytes were identified as shown in Figure S1A. Up to 10,000 cells of each cell population were sorted into low-retention 1.5-ml collection tubes (Thermo Fisher Scientific), containing 0.5 ml of a 1:1 solution of phosphate-buffered saline (PBS):FBS supplemented with ribonuclease inhibitor (1:100; Takara Bio). For some samples, directly after sorting, a small fraction of the T cell-Monocyte complexes sorted population was re-acquired on the same instrument, to confirm that the sort resulted in the physical separation of cells forming complexes.

### Single-cell RNA and TCR sequencing with 10X Genomics platform

Single-cell RNA sequencing was performed using the droplet-based 10X Genomics platform according to the manufacturer’s instructions. T cells and monocytes forming complexes and singlets were sorted as described in the cell sorting section. For ATB, we performed three experiment runs containing six samples each. For Dengue, we performed two experimental runs containing seven and eight samples, respectively. For each experiment run, PBMC samples were stained with a distinct hashtag oligonucleotide antibody as described in the flow cytometry section, to determine the sample origin for each cell after sequencing. Following cell sorting, cells were washed with ice-cold PBS, centrifuged for 10 min (600g at 4°C), and gently resuspended in ice-cold PBS supplemented with 0.04% ultrapure bovine serum albumin (Sigma-Aldrich). Cells were sorted by flow cytometry into low retention 1.5 ml collection tubes, containing 500 μl of PBS:FBS (1:1) supplemented with RNase inhibitor (1:100). After sorting, ice-cold PBS was added, cells were spun down, and single-cell libraries were prepared as per the manufacturer’s instructions (10X Genomics). Samples were processed using 10x 5’v2 chemistry as per the manufacturer’s recommendations. The library preparation was performed using Chromium Next GEM Single cell Standard 5V2 (Dual index) with feature Barcode technology kit and chromium Single Cell Human TCR Amplification Kit. Libraries were sequenced using the Illumina NovaSeq 6000 sequencing platform.

### Microscopy analysis of fixed T cell-monocyte complexes

For the visualization of CD3 polarization on T cell-monocyte complexes, thawed PBMC were incubated with live/dead Zombie UV (Biolegend, 1:1000 dilution) and 5 μl of FcR blocking reagent in 100 μl of PBS for 15min in the dark at room temperature. Cells were then washed with PBS supplemented with 10% FBS, resuspended in 100ul of PBS with 10% FBS and 2 μl of anti-human CD3 AF488 (clone UCHT1, Biolegend), 2 μl of anti-human CD14 BV480 (clone M5E2, BD Biosciences) and incubated for 20min in the dark at room temperature. Cells were washed twice with staining buffer (PBS containing 0.5% FBS and 2 mM EDTA, pH 8), resuspended in 100 μl Cyto-Fast Fix/Perm buffer (Biolegend), and incubated for 20min in the dark at room temperature. Cells were washed twice with Cyto-Fast Perm Wash buffer, resuspended in 0.5–1 mL of staining buffer, and kept at 4°C until sorting. Cell sorting was performed on a BD S6 cell sorter (BD Biosciences). From the live singlet cell gate, T cell-monocyte complexes, singlet T cells, and singlet monocytes were identified as CD3+CD14+, CD3+CD14- and CD3-CD14+ respectively, and sorted in staining buffer. After the sort, cells were centrifuged at 600g for a few minutes, resuspended in 100 μl staining buffer and each population was plated on a separate well of a µ-Slide 8 Well Glass Bottom chamber (Ibidi). Microscopy images were acquired using a 20x 0.8NA objective with Zeiss LSM880 confocal system. Laser lines (405, 488, 561, 633 nm) were directed to the sample with 405 nm and 488/561/633 nm main beam splitters. Imaging was set up in 4 tracks, detecting AF647 fluorescence in the Airyscan detector with 660 nm long-pass filter, AF568 fluorescence in Ch2 detector (577-629 nm), AF488 fluorescence in ChS1 detector (499-543 nm), BV421 fluorescence in Ch1 detector (412–456), and BV480 in ChS1 detector (499-543 nm). Pixel dwell time was 7.83 µs, unidirectional scanning was done with line sequential mode, and 1.09 Airy Unit pinhole size (for 488 nm excitation). Voxel size was set to 0.12 x 0.12 x 1.2 µm, and Z-stack spanning whole cells were recorded and maximum intensity projected. Images were analyzed with QuPath (version 0.5.0-rc2) (48).

### Single-cell RNA-seq data processing for the ATB dataset

The FASTQ files from the single-cell libraries were put into the 10X Genomics Cell Ranger function multi pipeline (v7.0.0) for alignment (to GRCh38 v2020-A, GENCODE v32/Ensembl 98), and demultiplexing (using the 3’ cell multiplexing pipeline). After this step, data were converted into demultiplexed outputs, and cells with zero or more than one positive sample barcode detected were discarded. A Seurat object was built with Cell Ranger outputs using R package Seurat (v4.9.9) (49) and R (v4.2.2). To remove low-quality and doublet cells, only cells with a percentage of mitochondrial genes lower than 8%, a total number of genes comprised between 1,000 and 4,500, and a total number of reads lower than 17,000 were retained. Next, to reduce the variance incurred by the diversity of TCR genes and their highly individual-specific expression pattern, which can potentially lead to the formation of individual-specific small clusters that do not represent biologically meaningful subsets, we aggregated raw counts of TCR genes into four subgroups: TCRA, TCRB, TCRG, and TCRD. The raw counts of individual TCR genes were removed from the count matrix, and the sums of raw counts of genes in each subgroup across individual cells were added to the count matrix. After TCR gene aggregation, an integration step was performed to correct for batch effect across three experimental runs. Total DO and SO cell numbers are indicated in Figure S1C. No doublet origin cells were recovered for one sample (participant TB10, diagnosis visit). Since this study aimed to compare doublet and singlet origin cells, this sample was excluded from the subsequent analysis. The Seurat object was split up into a list of three Seurat objects for each sequencing run. Each Seurat object was first normalized using version 1 of SCTransform function (50, 51). Next, function SelectIntegrationFeatures (setting nfeatures = 3000), PrepSCTIntegration, RunPCA, and FindIntegrationAnchors (setting normalization.method = “SCT”, and reduction = “rpca”) were run consecutively to rank top features and prepare the list of three Seurat objects for integration. This step was then followed by running the function IntegrateData (setting normalization.method = “SCT”, k.weight = 100) to integrate three runs of Seurat objects into one integrated Seurat object. Dimensionality reduction and clustering analysis were performed on integrated assay using the function RunPCA (setting dims = 1:30, k.param = 100) for dimensionality reduction, the function FindClusters (resolution = 0.6) for identifying clusters, and the function RunUMAP (setting dims = 1:30, metric= “cosine”) for visualization. T cells and monocytes clusters were annotated based on the expression of T cell or monocyte specific markers genes and split from the original Seurat Object into one T-cell and one monocyte Seurat object. The raw RNA counts of T cells from clusters 2, 4, 6, 7, 9, 10, 11, 12, and 13 were extracted for building the T cell object, and raw counts of monocytes from clusters 0, 1, 3, 5, 8 and 14 were extracted for building the monocyte object. The same integration steps (and parameters) as for the global Seurat object were performed on the T cell and monocyte Seurat objects to correct for batch effects across the three experimental runs. For dimensionality reduction, clustering, and visualization steps, the same functions and parameters were used except that the resolution was set to 0.8 and 0.3 for the T cell and the monocyte Seurat objects, respectively. To determine the resolution for identifying clusters, R package clustree (v0.5.1) (52) was used. Function NormalizeData and ScaleData were used to generate data slot and scale.data slot, respectively, in RNA assay for both subset Seurat objects for further downstream analysis. Top genes for each cluster were extracted using the FindAllMarkers function on SCT assay and data slot (setting min.pct = 0.25, logfc.threshold = 0.25, test.use = “wilcox”), selecting only the positive genes (adjusted p-value < 0.05).

### Single-cell RNA-seq data processing for the dengue dataset

The same steps were performed on the dengue dataset, from preprocessing steps using Cell Ranger to quality control, TCR genes aggregation, integration, normalization, dimensionality reduction, clustering, and visualization, with the following changes: i) during the quality control step, we retained cells with a percentage of mitochondrial genes lower than 15%, a total number of genes comprised between 200 and 5,000, and a total number of reads lower than 25,000; ii) for clustering, resolution was set to 0.7 based on clustree’s result.

### Single-cell RNA sequencing data analysis

For the ATB dataset, differential expression (DE) analyses were performed using the function FindMarkers on RNA assay and data slot using the MAST test (53) in 17 samples in total (Figure S1C for individual sample breakdown). In the analyses of DO versus SO T cells/monocytes, sample identifiers were used as a latent variable to account for inter-individual variability. For the comparison of doublet versus single origin T cells/monocytes within individual clusters, we did not control for inter-individual variability as the number of DO cells per sample per cluster were too small. For the dengue dataset, differential expression (DE) analyses were performed using the function FindMarkers on RNA assay and data slot using the MAST test (53) in 15 samples (Figure S4A for individual sample breakdown) without controlling for inter-individual variability, as too few samples had both doublet and SO cells retrieved. Genes were considered significant if their adjusted p-value was lower than 0.05 and their absolute log2 Fold Change was higher than 0.2. Gene scores were calculated by summing up the normalized counts (stored in the RNA assay data slot) of all genes in a given list for each cell. UMAP plots and dot expression plots were performed using the function DimPlot and DotPlot with R package Seurat (v4.9.9) in R (v4.2.2) (54). Heatmaps were performed using the function Heatmap with R package ComplexHeatmap (v2.15.4) (55). All the other graphic visualization figures, including volcano plots, dot plots, violin plots, and boxplots were generated using R package ggplot2 (v3.4.2), ggpubr (v0.6.6) and, ggsignif (v0.9.4).

### Single-cell TCR sequencing data analysis

TCR data was analyzed using R package scRepertoire (v1.8.0) (56). We first constructed a TCR genes table combining all Cell Ranger output files filtered_contig_annotations.csv for each sample using the function combineTCR. Next, we filtered cells that had both TCRA and TCRB genes detected, resulting in 24,025 cells across all 17 samples (Figure S1E for individual sample breakdown). For each sample, we performed random downsampling to the smallest sample size among the following four categories: DO at diagnosis, DO at end-of-treatment, SO at diagnosis and SO at end-of-treatment. This process yielded a total of 3,056 cells, whose clonotype information was then attached to the T-cell subset Seurat Object (setting cloneCall= “strict”, chain = “both”). Three clonotype groups were generated according to the relative proportion of cells expressing a given clonotype in all four categories. Clonotypes with cell counts less than 5 were labeled as small; clonotypes with cell counts between 5 and 20 were labeled as medium, and clonotypes with cell counts higher than 20 were labeled as large. Clonotype overlap between doublet and SO T cells was analyzed using the function compareClonotypes. TRAV, TRAJ, TRBV, and TRBJ gene usage was analyzed using function vizGenes. Finally, TCR CDR3B sequences were put into TCRMatch too (v1.1.2) (25) to identify its antigen specificity within the IEDB database (57) (setting -s the match score threshold to 0.9).

### Statistical analysis

Statistical analyses were performed using GraphPad Prism Software (version 10.2), or R (version 4.2.2). Paired datasets were compared using non-parametric Wilcoxon tests, while unpaired datasets were compared using non-parametric Mann-Whitney tests. P values less than 0.05 were considered significant and 2-tailed analyses were performed. Correction for multiple comparisons was performed with Bonferonni correction. Statistical significance of overlap between gene lists was calculated using the hypergeometric distribution test and considering all genes that were detected within T cells or Monocytes as the total number of genes (27,506 for T cells, and 28,365 for monocytes).

## Data availability

Flow cytometry data is available on the Immport portal under accession ID SDY2734, and single-cell RNA and TCR sequencing are available in GEO under accession number GSE273019.

## Acknowledgments

We thank the Flow cytometry, Sequencing, and Bioinformatics cores facilities at La Jolla Institute for Immunology for technical assistance with cell sorting, sequencing, and data analysis. This work was supported by grants from the National Institute of Allergy and Infectious Diseases of the National Institutes of Health under award numbers (U19-AI118626; AI62100; S10OD025052; S10OD021831; S10OD030417; S10RR027366) and the Conrad Prebys Foundation. The funders had no role in study design, data collection and analysis, decision to publish, or preparation of the manuscript.

## Authors contributions

JGB and BP conceived and designed the study. HH, CK, ZM, GS, PV, and JGB conducted the experiments. NK, AC, RT, and JGB performed data analysis. TJS, ADS, DW, AB, EH, AS, RT, and CLA provided clinical samples. NK and JGB led the data interpretation with input from BP, RT, and CLA. NK and JGB wrote the manuscript, and all authors edited the manuscript. All authors contributed to the article and approved the submitted version.

