## Supplementary material for "A novel method for characterizing cell-cell interactions at single-cell resolution reveals unique signatures in blood T cell-monocyte complexes during infection": Table S1, Figure S1, Figure S2, Figure S3, Figure S4: Biorxiv_TMDoublets_SuppMaterial.pdf

**Table S1**: **Active tuberculosis and dengue fever cohorts characteristics.**

| Donor ID | Cohort | Age | Gender | Race | Visits | Dengue Severity |
|---|---|---|---|---|---|---|
| TB1 | Active Tuberculosis | 22 | Female | More than 1 | Diagnosis; End-treatment | - |
| TB2 | Active Tuberculosis | 23 | Male | Black or AA | Diagnosis; End-treatment | - |
| TB3 | Active Tuberculosis | 50 | Female | More than 1 | Diagnosis; End-treatment | - |
| TB4 | Active Tuberculosis | 29 | Male | More than 1 | Diagnosis; End-treatment | - |
| TB5 | Active Tuberculosis | 26 | Female | More than 1 | End-treatment | - |
| TB6 | Active Tuberculosis | 35 | Male | More than 1 | Diagnosis | - |
| TB7 | Active Tuberculosis | 26 | Male | More than 1 | Diagnosis; End-treatment | - |
| TB8 | Active Tuberculosis | 46 | Female | More than 1 | Diagnosis; End-treatment | - |
| TB9 | Active Tuberculosis | 52 | Male | More than 1 | Diagnosis; End-treatment | - |
| TB10 | Active Tuberculosis | 29 | Female | Black or AA | Diagnosis; End-treatment | - |
| D1 | Dengue fever | Unknown | Male | Unknown | Acute; Convalescent | Non-Hemorraghic |
| D2 | Dengue fever | Unknown | Male | Unknown | Acute; Convalescent | Non-Hemorraghic |
| D3 | Dengue fever | Unknown | Male | Unknown | Acute; Convalescent | Non-Hemorraghic |
| D4 | Dengue fever | Unknown | Male | Unknown | Acute; Convalescent | Hemorraghic |
| D5 | Dengue fever | Unknown | Male | Unknown | Acute; Convalescent | Hemorraghic |
| D6 | Dengue fever | Unknown | Male | Unknown | Acute | Non-Hemorraghic |
| D7 | Dengue fever | Unknown | Male | Unknown | Acute | Non-Hemorraghic |
| D8 | Dengue fever | Unknown | Male | Unknown | Acute | Non-Hemorraghic |
| D9 | Dengue fever | Unknown | Male | Unknown | Convalescent | Non-Hemorraghic |
| D10 | Dengue fever | Unknown | Male | Unknown | Convalescent | Non-Hemorraghic |

A

B

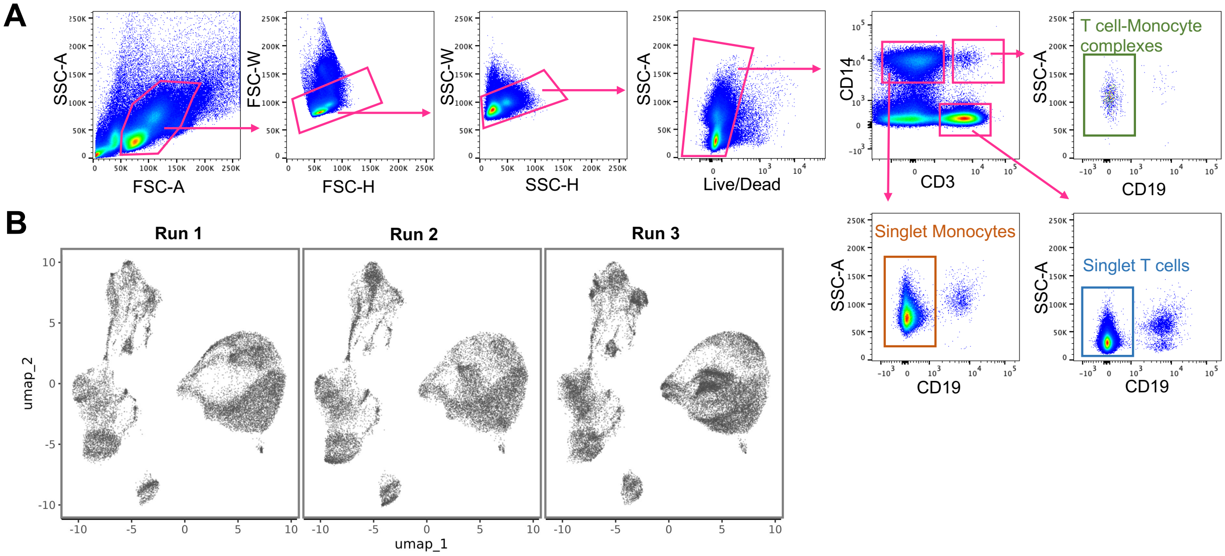


C

| Timepoint | Diagnosis | | | | End-Treatment | | | |
|---|---|---|---|---|---|---|---|---|
| Cell subset | T cells | | Monocytes | | T cells | | Monocytes | |
| Origin | Doublet | Singlet | Doublet | Singlet | Doublet | Singlet | Doublet | Singlet |
| TB1 | 69 | 1925 | 262 | 2033 | 75 | 1677 | 133 | 2004 |
| TB2 | 100 | 1671 | 118 | 1908 | 420 | 1046 | 499 | 1189 |
| TB3 | 128 | 1595 | 113 | 1865 | 397 | 1541 | 346 | 1686 |
| TB4 | 82 | 1575 | 530 | 1752 | 145 | 1633 | 246 | 1429 |
| TB5 | - | - | - | - | 129 | 2067 | 319 | 2445 |
| TB6 | 20 | 1818 | 1224 | 2370 | - | - | - | - |
| TB7 | 297 | 1540 | 534 | 1849 | 751 | 1715 | 984 | 2112 |
| TB8 | 64 | 1996 | 110 | 2424 | 112 | 1746 | 163 | 2154 |
| TB9 | 175 | 934 | 436 | 1296 | 231 | 961 | 516 | 1146 |
| TB10 | 0* | 1534* | 0* | 1510* | 90 | 1517 | 97 | 1608 |

*Sample excluded

D

| Timepoint | Diagnosis | | | | End-treatment | | | |
|---|---|---|---|---|---|---|---|---|
| Origin | Doublet | | Singlet | | Doublet | | Singlet | |
| TCRab | # cells with TCRab | % TCRab+ T cells | # cells with TCRab | % TCRab+ T cells | # cells with TCRab | % TCRab+ T cells | # cells with TCRab | % TCRab+ T cells |
| TB1 | 59 | 86% | 1558 | 81% | 62 | 83% | 1344 | 80% |
| TB2 | 76 | 76% | 1295 | 78% | 255 | 61% | 774 | 74% |
| TB3 | 110 | 86% | 1403 | 88% | 327 | 82% | 1308 | 85% |
| TB4 | 71 | 87% | 1329 | 84% | 127 | 88% | 1344 | 82% |
| TB5 | - | - | - | - | 104 | 81% | 1730 | 84% |
| TB6 | 13 | 65% | 1235 | 68% | - | - | - | - |
| TB7 | 220 | 74% | 1211 | 79% | 434 | 58% | 1160 | 68% |
| TB8 | 49 | 77% | 1687 | 85% | 92 | 82% | 1363 | 78% |
| TB9 | 127 | 73% | 769 | 82% | 190 | 82% | 812 | 85% |
| TB10 | 0* | NA* | 1268* | 82%* | 78 | 87% | 1309 | 86% |

*Sample excluded

**Figure S1: Post-sort and post-sequencing quality checks from the high-throughput single-cell analysis of T cells and monocytes forming complexes in human blood during active tuberculosis**. This figure is the supplement for Figure 1. A) Gating strategy to identify and sort T cell-Monocyte complexes, singlet T cells, and singlet monocytes from cryopreserved human PBMC using flow cytometry. B) UMAP representation of all cells (as depicted in Figure 1D), split by experimental run. C) Numbers of singlet and doublet origin T cells and monocytes obtained per sample after removing suspected doublets and low-quality cells. T cells and monocytes were identified based on the UMAP as shown in Figure 1D. D) Number and frequency of singlet and doublet origin T cells with both TCRα and TCRβ chains detected by TCRαβ sequencing obtained per sample.

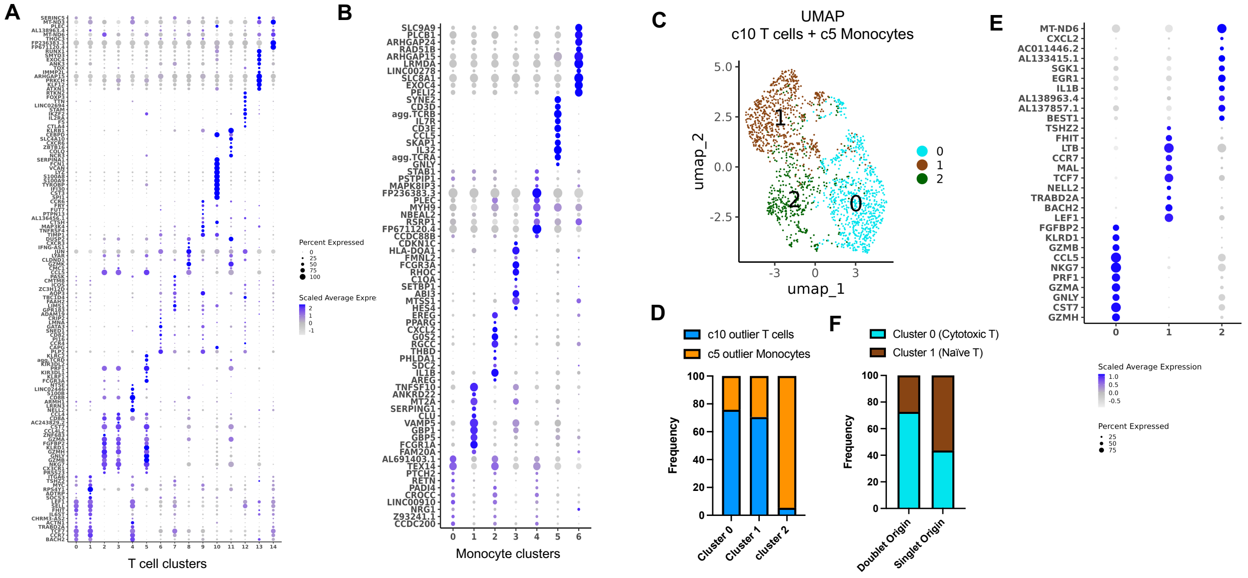


**Figure S2: Additional analyses in T cell and monocyte clusters enriched for cells of doublet origin**. This figure is the supplement for Figures 2 and 3. A) Dot plot representing the expression of the top 10 marker genes per cluster across all 15 T cell clusters identified in Figure 2E. B) Dot plot representing the expression of the top 10 marker genes per cluster across all seven monocyte clusters identified in Figure 3F. C) UMAP representation and clustering of the monocyte-like T cell cluster 10 (as identified in Figure 2E) combined with the cytotoxic T cell-like monocyte cluster 5 (as identified in Figure 3F). D) Frequency of T cells and monocytes within each cluster from the combined UMAP shown in C). E) Dot plot representing the expression of the top 10 marker genes per cluster across all three clusters identified in C). F) Frequency of cells classified in cluster 0 or cluster 1 (as identified in C) in doublet origin or singlet origin monocyte-like T cells.

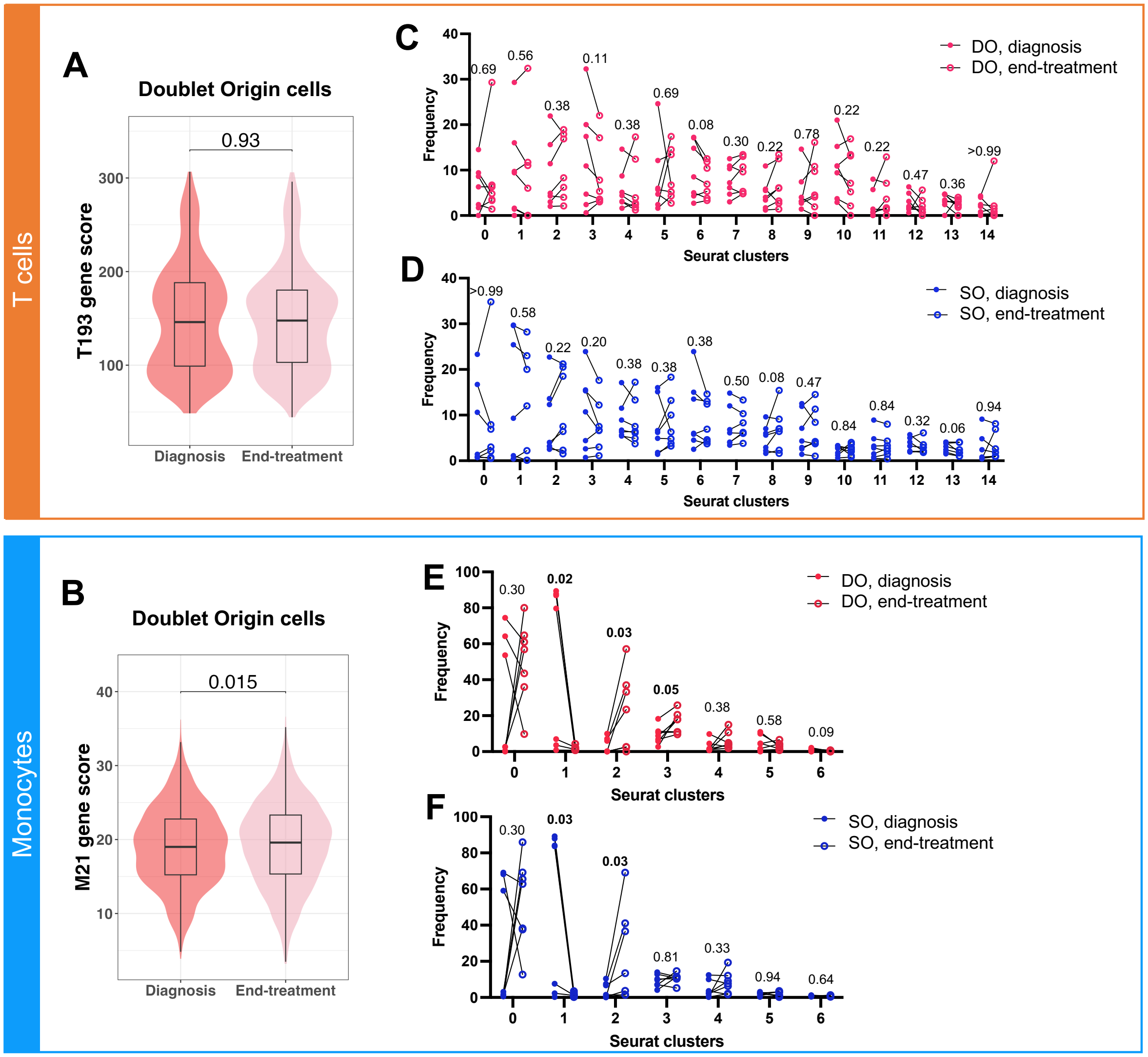


**Figure S3: Comparative analysis of the transcriptome of doublet origin T cells and monocytes at diagnosis versus end-treatment**. This figure is the supplement for Figure 4. A) Distribution of the T193 gene score in doublet origin T cells at diagnosis versus end-treatment. The T193 gene signature represents the sum of 193 genes significantly upregulated in doublet origin (DO) versus singlet origin (SO) T cells, as defined in Figure 2A. B) M21 gene score in doublet origin monocytes at diagnosis versus end-treatment. The M21 gene signature represents the sum of 21 genes significantly upregulated in DO versus SO monocytes, as defined in Figure 3A. In A-B), the lower, median and upper edges of the boxplots represent the 25th, 50th and 75th percentile; the length of upper and lower whiskers is 1.5 times the interquartile range. Non-parametric unpaired Mann-Whitney tests were used for comparison, and Bonferroni correction was performed to adjust the p-value. Differences in cell cluster composition between diagnosis and end-treatment paired samples in C) DO T cells, D) SO T cells, E) DO monocytes, and F) SO monocytes. T cell clusters were defined in Figure 2E and monocytes clusters were defined in Figure 3F. Non-parametric paired Wilcoxon tests were used for comparison.

**A**

| Sample ID | Participant ID | Timepoint | Doublet origin cells recovered | Singlet origin cells recovered |
|---|---|---|---|---|
| 1 | D1 | Acute | 0 | 1,323 |
| 2 | D2 | Acute | 0 | 797 |
| 3 | D3 | Acute | 219 | 169 |
| 4 | D4 | Acute | 125 | 113 |
| 5 | D5 | Acute | 101 | 89 |
| 6 | D6 | Acute | 188 | 0 |
| 7 | D7 | Acute | 196 | 195 |
| 8 | D8 | Acute | 268 | 223 |
| 9 | D1 | Convalescent | 0 | 163 |
| 10 | D2 | Convalescent | 801 | 0 |
| 11 | D3 | Convalescent | 147 | 112 |
| 12 | D4 | Convalescent | 33 | 0 |
| 13 | D5 | Convalescent | 16 | 0 |
| 14 | D9 | Convalescent | 0 | 151 |
| 15 | D10 | Convalescent | 340 | 0 |
| | | Total | 2,434 | 3,335 |

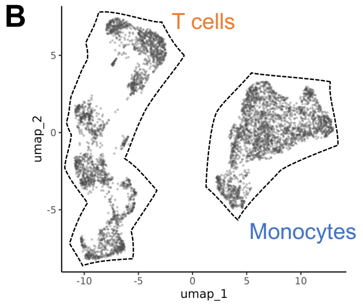


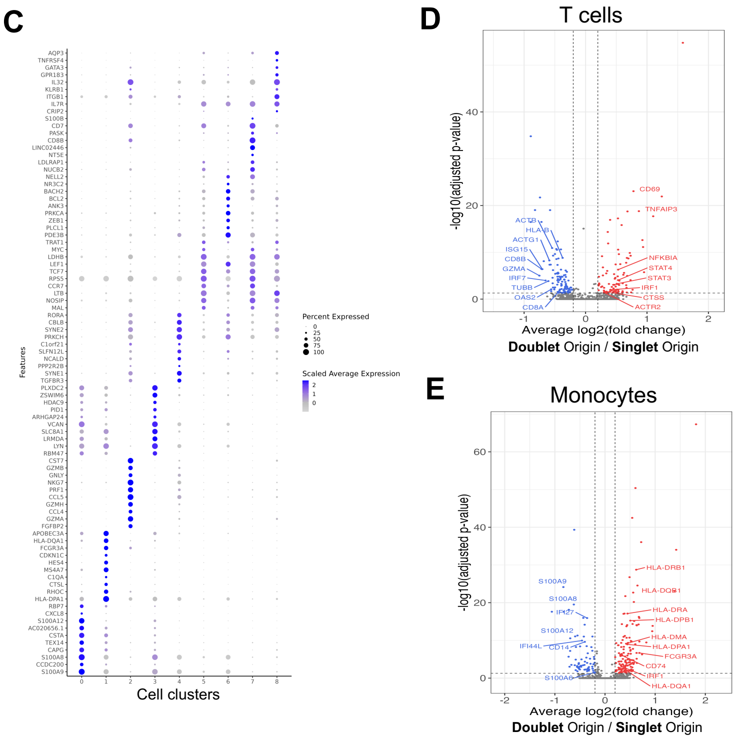


**Figure S4: Single-cell transcriptomic analysis of T cells and Monocytes forming complexes in human blood during dengue**. This figure is the supplement for Figure 6. A) Numbers of singlet and doublet origin cells after removal of suspected doublets and low-quality cells obtained per sample. B) UMAP and clustering of all cells. C) Dot plot representing the expression of the top 10 marker genes per cluster across all nine clusters identified in B). Volcano plots of differentially expressed genes (adjusted p-value with Bonferroni correction <0.05, average log2 fold change > 0.2) comparing D) T cells of doublet versus singlet origin, and E) monocytes of doublet versus singlet origin.
